# Density dependent survival drives variation in density dependent population growth of an insect pest

**DOI:** 10.1101/2023.11.21.568113

**Authors:** Shyamsunder Buddh, Sandeep Krishna, Deepa Agashe

**Affiliations:** National Centre for Biological Sciences, TIFR, Bengaluru, India

**Keywords:** Density dependence, demographic traits, *Tribolium castaneum*, competition, life history, flour beetle

## Abstract

Several ecological processes – from population dynamics to species co-existence – are driven by density dependence (DD) in population growth rate. Thus, to predict and manage ecological outcomes, we need a deep understanding of which factors and demographic traits drive variation in DD. In the insect pest *Tribolium castaneum*, we found large variation in DD across habitats but not across source populations. We modeled DD in population growth as the product of DD in fecundity and survival, experimentally estimating each parameter. Across habitats, survival parameters varied more than fecundity, including in simulations with varying parameter values. Thus, DD in survival drives variation in density-dependent population growth, and should evolve rapidly under strong density-dependent selection. Our general framework combining detailed experiments and simulations with a simple model can be used for other species to better understand the causes and consequences of density dependent population growth.

## INTRODUCTION

Changes in population size or density over time are often characteristic of species and habitats, and are important to understand their ecology and evolution. Decades of studies have firmly established that one of the main regulators of population density is density itself (Nicholson, 1957). This density dependence (“DD”) – the dependence of population growth rate on population density – can arise due to various mechanisms, e.g., predation (Hixon & Carr, 1997; Holbrook & Schmitt, 2002) or resource limitation (Amundsen et al., 2007). DD can manifest in either a negative or a positive slope for population growth vs. density (e.g., see Allee et al., 1949; Courchamp et al., 1999; Herrando-Pérez et al., 2012) with linear, concave or convex shapes (C. W. Fowler, 1981; Sibly et al., 2005). The slope and shape of DD have important consequences, including for population dynamics (reviewed in Dey & Joshi, 2018), spatial distribution and oviposition behaviour (Wetzel, 2014; Wetzel & Strong, 2015), the strength of inbreeding depression (Yun & Agrawal, 2014), population persistence under environmental perturbation (Drake, 2005; Vonesh & De la Cruz, 2002), and maintaining species coexistence (Chesson, 2000; Comita et al., 2010; Lamanna et al., 2016; Webb et al., 1999). With concave DD, a population’s growth rate will rapidly decline as a function of increasing density at low densities, but remain relatively unchanged across high densities. In contrast, with convex DD, the largest impact of increasing density is at higher population densities. Thus, the shape of DD can determine the impact of factors such as immigration or predation that alter population density, and the appropriate strategy for population conservation or control (Powell 1988). Hence, DD has significant population- and species-level ecological and evolutionary outcomes, and a deeper understanding of the variation in DD and mechanisms of density dependent population growth is useful to predict and understand these outcomes.

Prior work shows substantial interspecific (Sibly et al., 2005) and intraspecific variation in the slope and shape of DD, both across environments (Agrawal et al., 2004; Miller, 2007; Underwood & Rausher, 2000a) and genotypes (Marks, 1982; Zaiats et al., 2021). What demographic traits underlie this variation in DD? Empirical work shows that DD can act on different demographic traits over multiple life stages, such as adult survival and reproduction, and juvenile survival and development rate (e.g. Altwegg, 2003; Mueller, 1988). For example, Cayuela et al., (2019) found positive DD in adult survival but negative DD in fertility. Prior studies have modelled DD in per capita population growth as an outcome of DD in underlying demographic traits (Cayuela et al., 2019; Hellriegel, 2000; Miller, 2007; Mueller, 1988; Powell, 1988; Tung et al. 2019; Vonesh & De la Cruz, 2002). However, very few studies have directly quantified whether and how variation in DD in demographic traits gives rise to variation in DD in per capita population growth. This is important because understanding how specific traits contribute to population dynamics can help us make more precise predictions for how the population would change, which may help in predicting its evolutionary trajectory or in determining the efficacy of population management strategies. For example, prior work using matrix models showed that adult survival is more important than juvenile survival for population size (Caswell, 1989), leading to changes to conservation strategies (reviewed in Benton & Grant, 1999). However, matrix models primarily deal with populations that have density-independent dynamics, and that are therefore either at equilibrium or changing monotonically in size. Hence, this approach is not useful for populations that fluctuate over a range of densities (e.g. Stiling, 1988), such as density-dependent populations (Dornelas et al., 2019; McGill, 2020) or populations not at equilibrium, e.g., fast growing, dispersal-limited species such as insects on islands. However, some studies have used matrix models to study density dependent population growth using DD in demographic traits (e.g. Benton & Grant, 1999), but this work is also limited because DD is estimated from field observations (but see Dennis et al., 1995; Grant & Benton, 2003), which is notoriously unreliable and biased (Detto et al., 2019; Knape, 2008; Knape & de Valpine, 2012). Notably, several studies (using life table response experiments; LTRE) have asked which life stages are most affected by changes in density and play the largest role in driving subsequent changes in population growth rate (e.g. Bassar et al., 2013; N. L. Fowler et al., 2006; Reznick et al., 2012). However, even these studies have not addressed how DD in multiple demographic traits gives rise to variation in the slope and shape of DD within a species, e.g., across different habitats or populations. To summarize, though it is important to identify demographic traits with the largest impact on variation in density-dependent population dynamics, there is no systematic study addressing this problem.

We conducted experiments on the red flour beetle *Tribolium castaneum*, a global generalist pest of stored grain products (Sokoloff, 1972) that shows DD in fecundity and survival (Johnson et al., 2022; Khan et al., 2018; Ravi Kumar et al., 2023; Sonleitner & Guthrie, 1991). Flour beetles have multiple life stages (egg, larva, pupa, adult) which can all be counted, enabling the partitioning of density dependence in population growth rate into density dependence in life-stage specific demographic traits. They also show cannibalistic behaviour (Sokoloff, 1972), but the opportunity for cannibalism was restricted in our experiment by the use of discrete generations. We first quantified variation in DD across habitats (four different grain flours) and source populations (ten populations collected from different parts of India and maintained in the laboratory on wheat flour). Finding significant variation across habitats, we then conducted further experiments to partition DD in population growth into DD in fecundity and survival. We built a discrete-time-step population dynamics model for each habitat by fitting difference equations to the demographic data, and successfully reconstructed DD in population growth rate from DD in fecundity and survival. Using simulations, we showed that density-dependent juvenile survival, rather than density-independent juvenile survival or density-dependent or -independent fecundity, has the largest impact on variation in DD in population growth rate in *T. castaneum*.

## MATERIALS AND METHODS

### Experiment 1: Quantifying variation in density dependent population growth across source populations and habitats

To test how DD varies across habitats and source populations, we used a full factorial design with habitat, source population and density as three factors. For the habitat treatment, we used four flours: whole wheat, corn (*Zea mays*), finger millet (*Eleusine coracana*), and sorghum (*Sorghum bicolor*). All flour was kept at -80 °C for at least four hours to kill any pests, and thawed for at least eight hours before use. For the population treatment, we used 10 source populations collected across India and maintained in the laboratory for 5-10 years. Population stocks were kept on an 8 week discrete generation cycle with at least 500 adults used to set up each generation. We replaced the flour and discarded dead individuals and larval moults at least once every generation. All stock populations and experimental boxes were kept at 34°C in a dark incubator. The sex ratio of beetles is usually balanced in stock populations, so we randomly sampled individuals from stock boxes of each source population to set up the experiment, conducted in 2018-2019. For the density treatment, we used ten founding densities of adults (adults/g flour): 0.6, 0.9, 1.2, 1.5, 2, 2.8, 3.6, 4.4, 6, and 7, and provided 50g of flour in each experimental box (leading to 30, 45, 60, 75, 100, 140, 180, 220, 300 and 350 adults in 50g flour).

We randomly assigned the collected adults across habitat and density treatments, and let them lay eggs for one week (Figure 1A). Then we removed the adults, let the offspring develop in the same flour for 3 weeks (in wheat or sorghum) or 4 weeks (in corn or finger millet), and then counted total offspring (larvae + pupae + adults), terminating the experiment. The extra week in the latter case was based on our prior observation that development time is longer in corn and finger millet.

**Figure 1.**
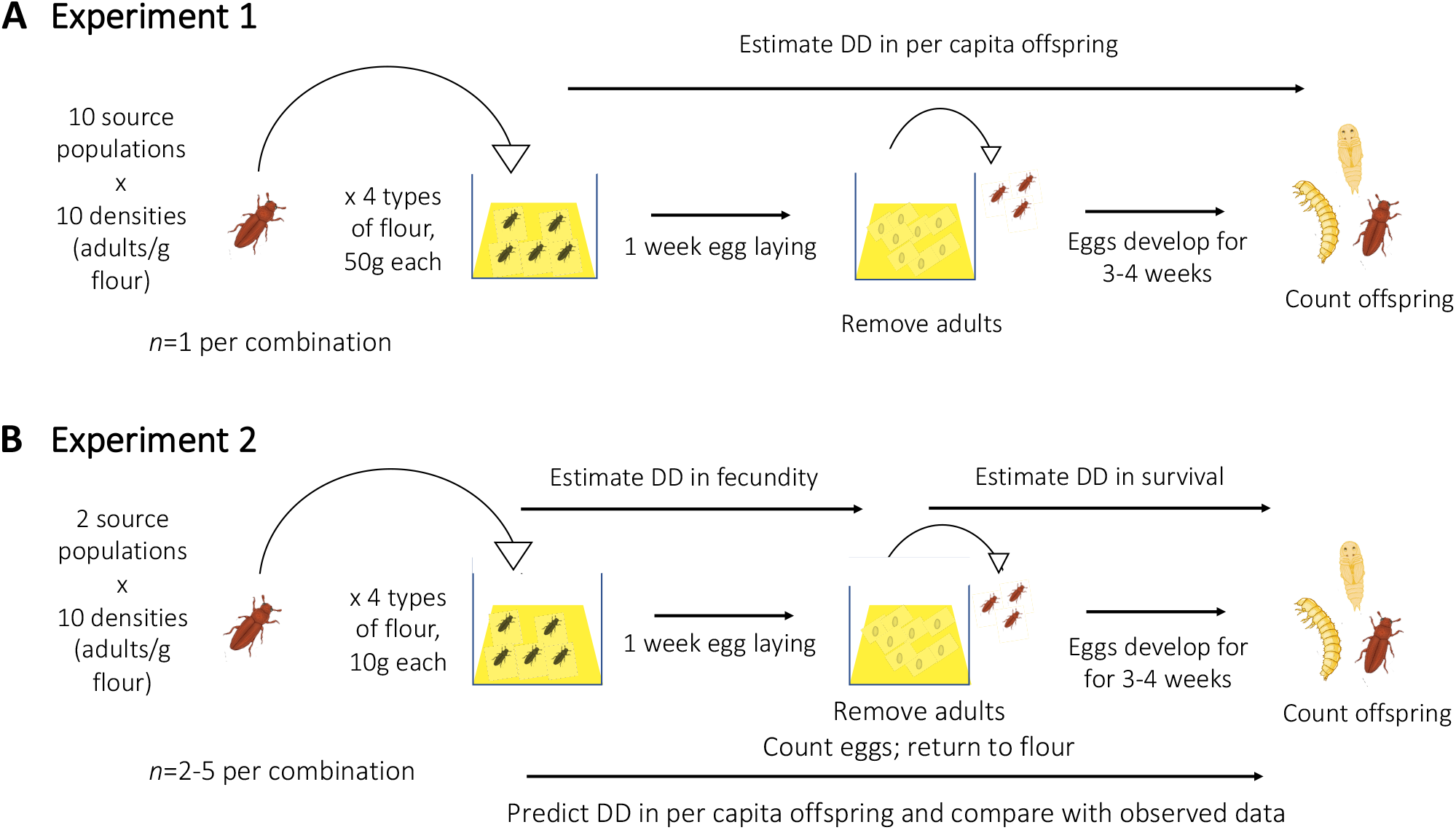
Schematic describing the two main experiments. (A) Experiment 1 was a fully factorial experiment with ten densities (0.6, 0.9, 1.2, 1.5, 2, 2.8, 3.6, 4.4, 6, & 7 adults/g flour) generated with adults from 10 distinct source populations, set up in four habitats. Here, we counted total developed offspring (larvae, pupae and adults) for each replicate. (B) Experiment 2 was similar but scaled down spatially, with fewer adults and only two populations, set up at identical densities with more replication. Here, we separately estimated fecundity and survival for each replicate. See Methods for further details.

We conducted all the data analysis and simulations described below using R Statistical Software (v4.2.0, R Core Team, 2022) in R Studio (v2022.12.0.353, Posit team, 2022), and used the package ‘ggplot2’ (Wickham, 2016), to make the figures.

We calculated per capita offspring = total offspring/founding no. of adults. Next, we asked how the slope and shape of DD, i.e., the relationship of per capita offspring and founding density (i.e., founding adults/g, henceforth density), changes across habitats and populations. To first determine the best model that fit data from each habitat and source population combination, we fit a linear model (per capita offspring = a*density+b), a nonlinear exponential model (y=a*e^b*density^), and a different nonlinear model (per capita offspring = a*density^b+c*density^), using ordinary least squares (for the linear model) and nonlinear least squares (for the nonlinear models). The second nonlinear model had significantly lower AIC values than the other two models (see dataset: Buddh et al., 2023), so this was our model of choice for nonlinear density dependence. For comparison, we then fit simplest linear model as a straight line only between the lowest and highest density points of the nonlinear model. We used the slope of this linear fit as the overall slope of density dependence. To quantify shape, we calculated the average value of the absolute difference between the y axis values predicted by the two models (nonlinear vs. linear), across ∼600 evenly spaced values along the range of densities represented on the x axis (Figure 1E). We quantified the shape this way because we wanted to keep measures of shape consistent across analyses, and because this provides a way to quantify the shape of a curve even if one doesn’t know the exact parameter that controls shape. We extracted the shape and slope for all population-habitat combinations, and analysed these data using an ANOVA with slope∼population+habitat and shape∼population+habitat. We calculated the effect size of each independent variable as the sum of squares of treatment/total sum of squares.

### Experiment 2: Measuring DD in demographic traits underlying DD population growth

To identify which demographic traits drive variation in DD in population growth, we set up a second fully factorial experiment in 2021-2022 with ten densities, four habitats and only two populations. Since we did not find a strong population effect in the previous experiment; we used two randomly chosen populations (7 and 12). To facilitate egg counting, we conducted a scaled-down version of the previous experiment, with 5 times less flour and 5 times fewer founding adults. Thus, founding density was consistent with the previous experiment, but the flour, adult number, and experimental containers were smaller (10g flour per experiment, with founding adult number ranging from 6 to 70). To account for increased variation at the lower scale, we increased the number of replicates at each density (Figure 1B). At the end of the one-week egg laying, we removed the adults, counted all eggs in each container, returned them to the same flour and let them develop for 3-4 weeks.

We estimated the total eggs laid in each container by multiplying the number of eggs counted after 1 week of oviposition by 3.5. This is because eggs hatch in ∼48h, eggs laid in the first 5 days would not be counted. Hence, we extrapolated to 7 days by multiplying the number of eggs laid in 2 days with 3.5. We calculated per capita fecundity = total eggs/number of adults and proportion survival = total offspring/total eggs. We then fit the following models (using nonlinear least squares) to a set of relationships that, when multiplied together, would generate per capita offspring as a function of density.

Per capita fecundity = *a x* density*^b^*

Total fecundity = per capita fecundity x density

Proportion survival = (1+*alpha* x (total fecundity/*K*))/(1+(total fecundity/*K*))

Per capita offspring = per capita fecundity x proportion survival

*a* is a parameter that determines per capita fecundity at low density. The other three parameters indicate the nonlinear rate of decline of per capita fecundity with adult density (*b*), the asymptotic survival approached at high egg densities (*alpha*), and the egg density required to reduce the proportion survival from 1 to *alpha* (*K*). We used egg density as a proxy for the strength of larval competition during development. We fit these models to the experimental data for density-dependent survival and fecundity (measured separately), and compared the predicted per capita offspring values with the observed values. The chosen nonlinear models for fecundity and survival had slightly lower AIC values compared to a linear model and an exponential model (see dataset: Buddh et al., 2023).

Note that the values of density used in this model range from 6 to 70. This is the density of adults per 10 g flour. In Figure 3, where these results are described, the x axis for adult density ranges from 0.6 to 7 i.e., the density of adults per gram flour – this is done just for visual purposes, to keep the x axis consistent between Figure 2 and Figure 3.

**Figure 2.**
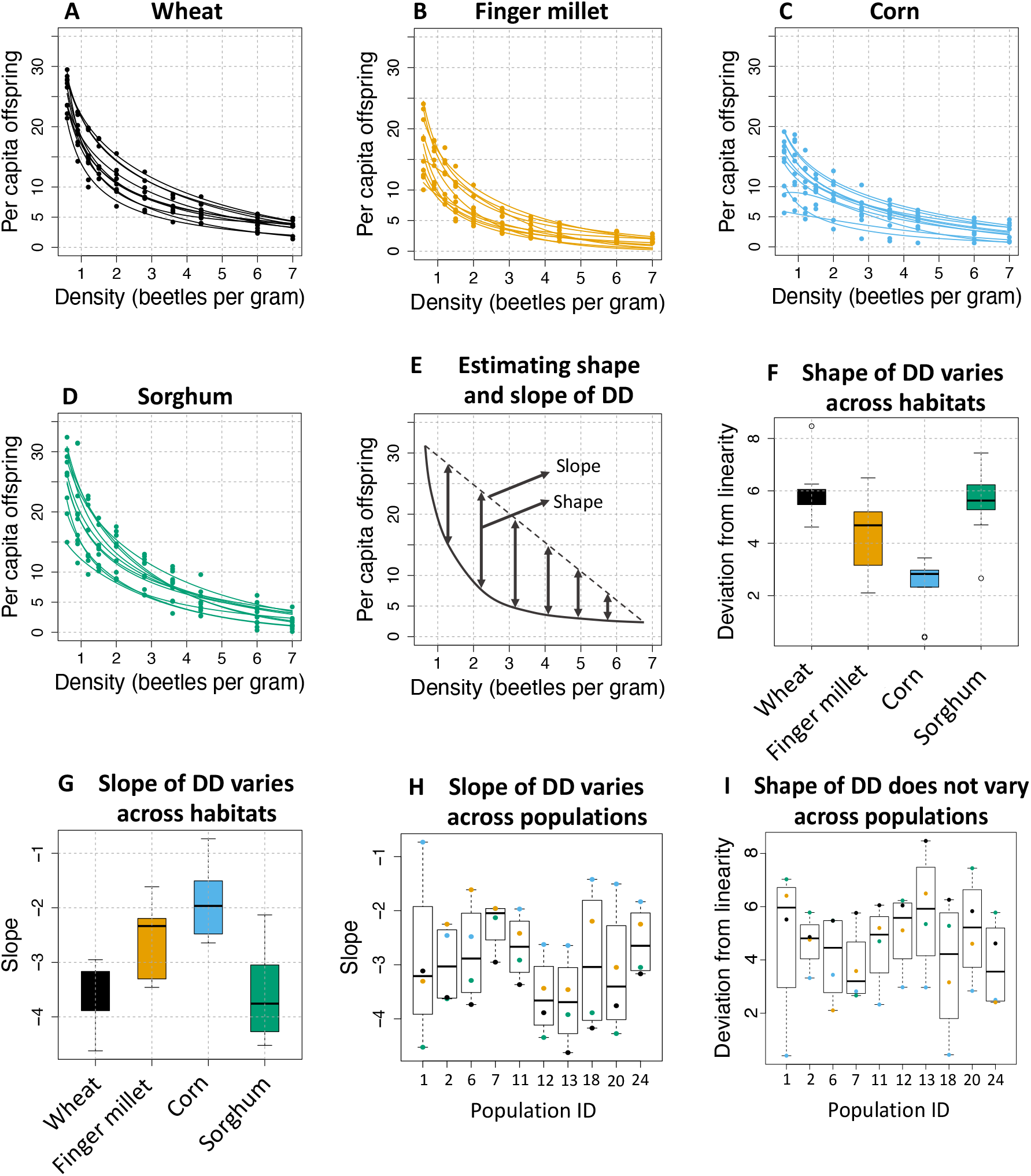
The slope of DD in population growth rate varies across populations and habitats, but the shape of DD in population growth rate varies only across habitats. The figure shows data from Experiment 1. (A–D) Per capita offspring (total offspring / number of founding adults) as a function of the density of founding adults in each habitat. Each point shows data for one replicate, and lines indicate best fit nonlinear models (of the form y=a*x^b+c*x^) for each source population (n=1/density/population/habitat). (E) Schematic showing how we quantified the slope and shape of DD in per capita population growth rate for each source population in each habitat. The slope was calculated by fitting a linear model to the data at lowest and highest density, and the shape was estimated as deviation from linearity, using the average value of the absolute difference between the fitted linear and non-linear models for ∼641 densities sampled across the x axis. (F, I) The shape of DD across habitats or populations (ANOVA, p_habitat_ = 2.2*10^-6^, p_population_ = 0.27). (G, H) The slope of DD in population growth rate across habitats or source populations (ANOVA, p_habitat_ = 2.1*10^-7^, p_population_ = 0.034).

**Figure 3.**
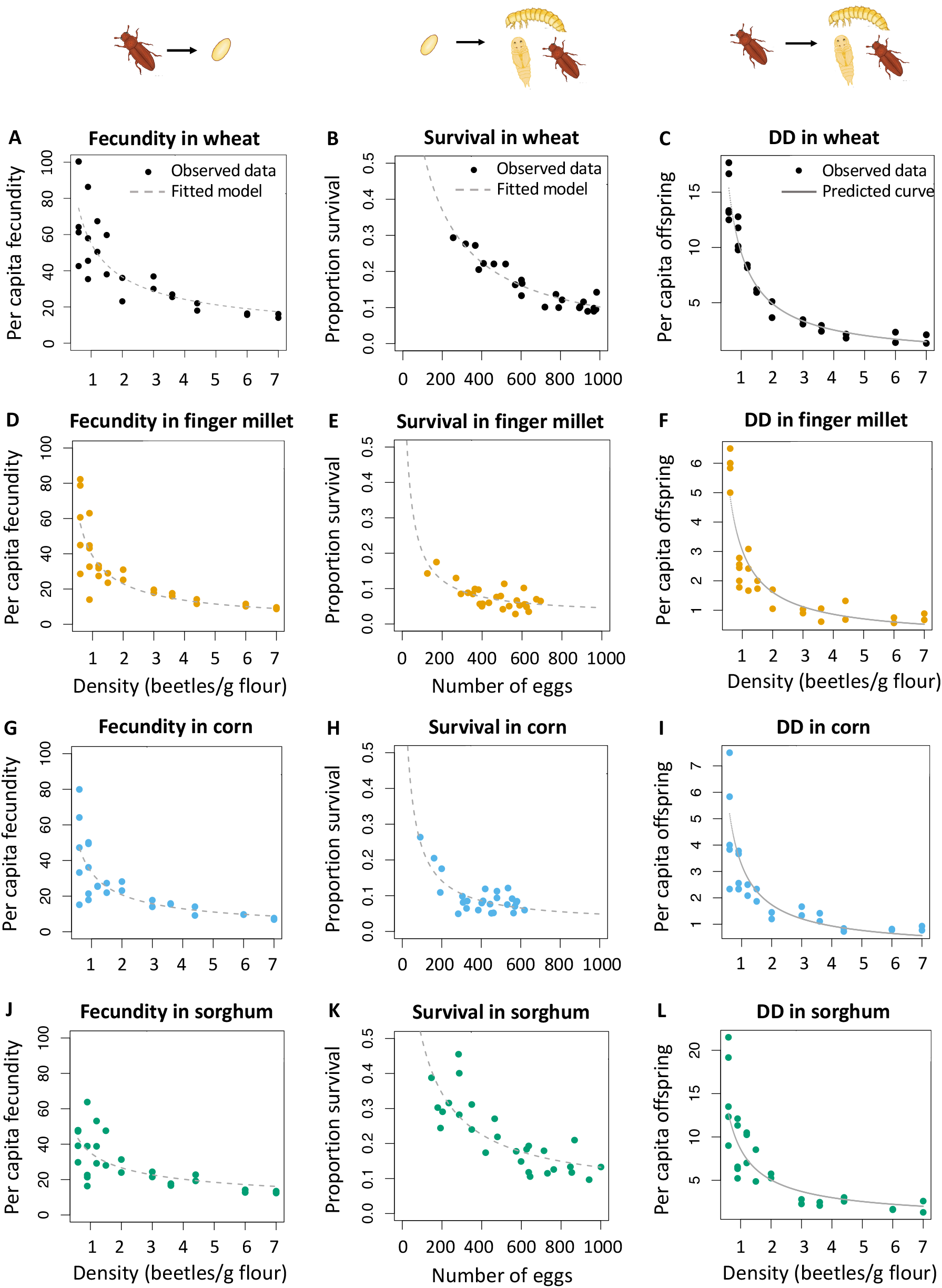
DD in fecundity and survival accurately predicts DD in per capita offspring. The figure shows data for population 7 measured in Experiment 2; data for population 12 are shown in Figure S2. (A–C) Density dependence in per capita fecundity (as a function of adult density), in survival (as a function of number of eggs), and in per capita offspring (as a function of adult density) in wheat flour. Each point represents experimentally measured data for a single replicate (n = 2–5 per density per habitat). In panels A and B, dashed lines indicate models fitted to the data using nonlinear least squares. In some cases, model fits extend beyond the range of observed data; the extrapolated fits were necessary for simulations shown in Figure 4. In panel C, the solid line shows the predicted density dependence in per capita offspring derived from multiplying the equations for models shown in panels A and B. Similarly, panels D–F show results for finger millet, G–I show results for corn, and J–L show results for sorghum. Model equations and estimated parameter values are shown in Table 1.

**Table 1.** Estimated parameter values and 95% confidence intervals (CI) for density-dependent fecundity and survival for population 7 in each of the four habitats (data shown in Figure 3). W = Wheat flour, C = Corn flour, FM = Finger millet flour, S = Sorghum flour. Per capita fecundity = *a*\*density*^b^.* Proportion survival = (1+*alpha*\*(total eggs/*K*))/(1+(total eggs/*K*)). Parameter values for population 12, and associated experimental data, are shown in Table S1 and Figure S2.

| Habitat | Fecundity $a$<br>(95%CI) | Fecundity $b$<br>(95% CI) | Survival $\alpha$<br>(95% CI) | Survival $K$<br>(95% CI) |
| --- | --- | --- | --- | --- |
| W | 215.8 (113.6) | -0.59 (0.21) | -0.006 (0.032) | 119.2 (23.9) |
| C | 160.3 (138.6) | -0.68 (0.33) | 0.022 (0.027) | 27.8 (9.4) |
| FM | 220.1 (159.7) | -0.75 (0.29) | 0.024 (0.022) | 22.7 (9.0) |
| S | 88.6 (45.3) | -0.40 (0.20) | 0.056 (0.071) | 87.5 (37.9) |

### Simulations to test the impact of survival and fecundity on the slope and shape of DD

To test the impact of variation in the model parameter values (described above) on variation in DD, we conducted simulations. Recall that for each habitat, the model fits involve 4 different parameters. We first fixed the value of 3 parameters for a given habitat, and varied the fourth parameter across the range of experimentally measured values across all four habitats, with a sampling frequency sufficient to obtain a large range of simulated values for each parameter (at least 60 intermediate values, and up to 372 intermediate values, depending on the parameter; Figure S1). Across these combinations of parameters values, we quantified the range of the slope and shape of the generated DD curve. The larger the range, the larger the impact of that parameter on DD in population growth. If the predicted curve dipped below 0 – which is not biologically relevant since it indicates negative per capita offspring – we removed all points following the first negative value of per capita offspring. We then repeated the simulations using parameter values for each habitat in turn. In total, we conducted 3104 simulations for population 7 and 2408 simulations for population 12.

The results of the above simulations could be influenced by differences in the range of values of each parameter across habitats. For instance, the values of parameter survival *K* range from 27.8 to 119.2 across habitats, whereas values of parameter fecundity *b* vary much less, from -0.4 to -0.75 (Table 1; also see Table S1). Therefore, we conducted a second set of simulations where we varied each parameter proportionally. To do this, for each habitat, we simulated a 1% change in one parameter value while keeping the other three parameter values fixed. Thus, we conducted 16 simulations for all combinations of parameter values for each habitat. As above, we calculated the difference in the slope and shape of DD in population growth rate with the new parameter value vs. the original experimentally measured value. We conducted both sets of simulations above using data from experiment 2, collected for two populations (population 7 data and parameters are shown in Figure 3 and Table 1 respectively; for population 12, see Figure S2 and Table S1).

## RESULTS

### Compared to source population, habitat has a larger effect on density dependent population growth

We first described the nature of DD (density dependence) in per capita offspring (our measure of population growth rate) for 10 source populations in four habitats across 10 densities (Experiment 1, Figure 1A; Figures 2A–D), and quantified the slope and shape of DD (Figure 2E). While the slope varied significantly across habitats (ANOVA; p=2.1*10^-7^, effect size = 0.57; Figure 2G) and source populations (p=0.034, effect size = 0.19; Figure 2H), the shape varied only with habitat (p= 2.2*10^-6^, effect size = 0.57; Figure 2F) and population did not have a significant effect (p=0.27; Figure 2I) (effect size was measured as the sum of squares for a treatment/total sum of squares). The variation in slope across populations was primarily driven by the difference between populations 13 and 7. The shape of DD was most nonlinear in wheat and sorghum, and most linear in corn, whereas the slope of DD was most negative in wheat and sorghum and least negative in corn. These results were broadly corroborated in a separate experiment with two adult densities, four habitats and four populations, where the slope of DD varied significantly across habitats but not populations (Experiment 3, Supplementary Methods; Table S2; Figure S3).

### DD in survival has a larger impact than fecundity on density dependent population growth rate

Next, in an experiment with two populations (populations 7 & 12; Experiment 2, Figure 1B), we measured DD in two key demographic traits – fecundity and survival – and asked whether their product accurately predicted DD in per capita offspring measured in the same experiment. We fit the following equation to the relationship between per capita fecundity and density in each habitat: per capita fecundity=a*density^b^, where *a* is the maximum per capita fecundity at low density (density-independent fecundity) and *b* is the nonlinear rate of decrease in fecundity with density (Figure 3 A, D, G, J, showing data for population 7; for population 12, see Figure S2). We fit the following equation for survival: proportion survival=(1+alpha*(total eggs/K))/(1+(total eggs/K)), where *alpha* is the proportion survival reached asymptotically as the number of total eggs increases, and *K* is the value of total eggs at which proportion survival reaches the midpoint between 1 and *alpha* (Figure 3 B, E, H, K). Then, we multiplied the two equations to predict the per capita offspring as a function of adult density. We found that in each habitat, the predictions matched observed values of per capita offspring very well (Figure 3 C, F, I, L; estimated parameter values are given in Table 1; see Figure S2 and Table S1 for results with population 12). Thus, our simple model accurately captured critical features of DD in demographic traits that ultimately drive DD in population growth.

Using this model, we tested whether DD in population growth was more sensitive to DD in fecundity or DD in survival. Parameter values for DD in survival showed more variation across habitats than parameter values for DD in fecundity (Table 1), suggesting a larger impact of density-dependent survival. To explicitly test this, we simulated variation in each parameter value in turn for the set of parameter values estimated for each habitat (see Methods; Figure S1), and calculated the range of resulting values of the slope and shape of DD in population growth. In both populations, the slope and the shape of DD in population growth rate were most sensitive to survival parameter *K*, followed by fecundity parameter *a* (Figure 4, Figure S4A–B). In contrast, fecundity parameter *b* had the weakest impact of all. Thus, for fecundity – where we could separate density independent and dependent effects (captured by parameters *a* and *b* respectively) – density independent effects were stronger. For survival, we could not explicitly evaluate density independent vs. dependent effects because in our model, density-independent survival (i.e., the intercept) is fixed at 1, and both *alpha* and *K* parameters together determine the nature of DD. In independent experiments, we verified that the intercept value for survival (i.e., survival for isolated eggs) is indeed close to 1 for all habitats (Experiment 4; survival ranges from 0.83–0.96 across habitats, Supplementary Data) and does not differ significantly across habitats (Table S3). We also note that isolated eggs are unlikely to be biologically relevant for *T. castaneum* since adult females continuously lay eggs, and population size is high given the high intrinsic growth rate of this species.

**Figure 4.**
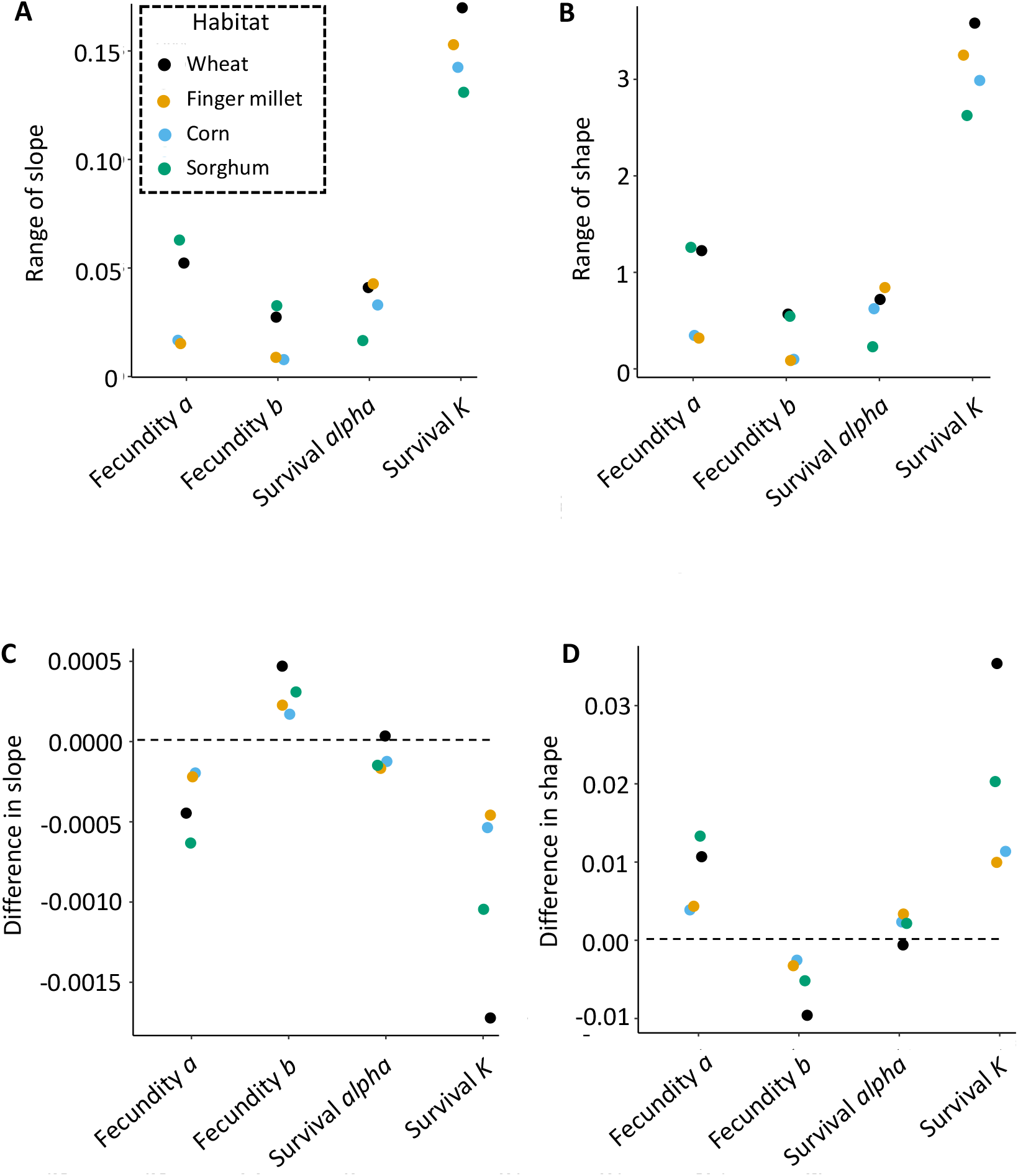
The slope and shape of DD in per capita offspring are most sensitive to survival parameter *K*. The plots show results of simulations to test the effect of varying each parameter value in our models describing DD in fecundity and survival for population 7 (Figure 3, Table 1). (A–B) For each habitat we varied the value of one parameter in turn (across the full range of values observed across habitats), and kept the other parameters constant. Plots show the range of the resulting values of the slope and shape of DD in per capita offspring, predicted as described in Figure 3 (across 3104 simulations). (C–D) Here, we again simulated variation in a single parameter value for a given habitat, but only increased the parameter value by 1% of the observed value. Points are slightly jittered about the x axis for clarity. Further details of the simulations are given in the Methods section and in Figure S1.

The differential impact of parameters on DD in population growth could reflect differences in the observed range of values of each parameter across habitats (e.g., compare the ranges of survival *K* with the range of fecundity *b* in Table 1 and Table S1, for populations 7 and 12 respectively). To evaluate this possibility, we carried out a second set of simulations, where we varied parameter values proportionally (instead of varying absolute parameter values as described above) by 1%, 5%, 20%, 50% or 100% of the original value. For population 7, proportional changes in parameter *K* again had the largest impact on the experimentally measured slope and shape of DD in population growth rate, followed by fecundity parameter *alpha* (Figure 4C–D; Figures S5 and S6). These results are consistent with the outcomes of the previous simulations, although in this case survival parameter *alpha* (rather than fecundity parameter *b*) had the weakest impact. For population 12, proportional changes (of 1%, 5% and 20%) in both survival parameter *K* and fecundity parameter *a* had equal and strong impacts (Figure S4C–D, Figure S7). Thus, for population 12, the larger effect of survival K appears to be driven by the wider range of observed parameter values across habitats, rather than inherently stronger impacts of survival compared to fecundity. Together, our simulations suggest that DD in survival has the largest effect on DD in population growth rate, and that density-independent fecundity has a larger impact than density-dependent fecundity.

## DISCUSSION

Understanding the processes driving population growth and dynamics has been a key goal of ecological research for over a century. Despite a large body of work with many species, explicit measurements of factors driving variation in density-dependent population growth rate, and the underlying mechanisms, remain scarce. Here, combining both experimental and modelling approaches for the red flour beetle, we first showed that DD varies most strongly across habitats, and less so across source populations. We then tested the impact of DD in two key demographic rates, finding that DD in survival has a larger effect on DD in population growth. Importantly, these results were robust across source populations and parameter values. Our work addresses a long-standing gap in population ecology and leads to several new insights and directions for further work, discussed below.

Our first main result is the strong effect of habitat on both the slope and the shape of DD. This is consistent with previous work on *T. castaneum*. For instance, a recent study reported distinct slopes and shapes of density dependent offspring production across different diets, possibly due to variation in dietary protein content (Đukić et al., 2022). Other insect species also show variation in DD across environments, e.g., phytophagous insects developing on different host plants (Agrawal, 2004; Miller, 2007; Underwood & Rausher, 2000b; Wetzel, 2014). Variation in DD should ultimately arise from variation in DD in specific demographic traits that underlie important life history parameters that determine population growth. Despite this, there is very little work on the processes that underlie the large variation in density dependent population growth across habitats. An interesting example is a field experiment showing that density-dependent population growth of cactus bugs is mediated by host plant quality, via its impact on juvenile bug survival (Miller, 2007). These results are strikingly similar to what we observed with flour beetles. For population 7, survival parameter *K* (which is the density required to kill approximately half the population, indicating the tolerance of juvenile survival to juvenile density) most affected the slope and shape of DD (Figure 4, S5, S6), both when considering the overall observed range of parameter variation as well as proportional variation in each parameter. On the other hand, for population 12 (Figure S4, S7), although survival parameter *K* still had the largest effect when considering the overall observed range of variation, proportional variation in survival *K* and fecundity *a* had equally strong effects on the slope and shape of DD. Overall, this suggests that variation in survival parameter *K* primarily drives variation in *T. castaneum*’s density dependence across habitats. In one of two populations, this effect is due to the larger empirical variation in this parameter, since its intrinsic impact (measured from varying the parameter proportionally) is the same as that of fecundity parameter *a*. However, the fecundity parameter is more constant across habitats while survival varies much more. It remains to be seen whether DD in survival is often the primary demographic trait that drives negative DD across species.

In contrast to the strong habitat effect, we saw only a weak impact of source population on the slope and shape of DD, despite substantial geographic distance, phenotypic variation, and adaptive trajectories across our source populations (Ravi Kumar, 2020). Some studies have reported variation in DD across genotypes (Marks, 1982; Zaiats et al., 2021), indicating genetic variation for density dependence. The lack of source population effects in our case may reflect low genomic differentiation across these populations (Shivansh Singhal, pers. Comm.), or loss of genetic variability during adaptation to laboratory conditions (Hoffmann & Ross, 2018) over 5-10 years after collection from the grain warehouses. Specifically, our populations may have very little variation in survival parameter *K*, which is the most critical parameter driving variation in DD in population growth. Finally, we considered that we may not have sufficient replication to detect a population effect. A power analysis indicates that in our experiment 1, the probability of detecting a moderate source population effect size of 0.4 with a p-value of 0.05 is 0.28, which is low. However, in another experiment with greater replication (Experiment 3, SI methods; n = 3 replicates for each habitat), we did not find significant variation in DD in per capita offspring across source populations, and only found a significant habitat effect (Table S2; Figure S3). This suggests that population level variation in DD is truly weak – perhaps due to less variation in survival parameter *K* – potentially constraining evolutionary change in DD.

Why is DD in population growth more sensitive to survival rather than fecundity? One way to approach this question is to ask whether this difference is evolutionarily meaningful. However, there has been little analysis of how sensitivity to DD in demographic traits should evolve, though prior work has addressed how density-independent demographic traits may evolve given stronger or weaker DD in population growth (Grant, 1998). Alternatively, our results may reflect the structure of our model, where survival follows fecundity. Alternative model formulations might start with survival, followed by fecundity (e.g., in an egg-to-egg recursive model), or involve both survival and fecundity feeding into each other across generations, e.g., when predicting multi-generation population dynamics. In our model, DD in survival may also compensate for variation in fecundity, potentially reducing the impact of density-dependent fecundity. For example, (Vonesh & De la Cruz, 2002) found that strong larval density dependence reduced the impact of variation in egg densities on adult population size, even creating a negative relationship in some cases. Finally, we note that in several organisms with density-independent population dynamics, survival has a larger role than fecundity in driving population growth rate (e.g. Pfister, 1998; Sæther & Bakke, 2000), though these studies also consider adult survival (which is not included in our model since the flour beetle adult lifespan is much longer than generation time). This may suggest a more general trend of greater importance of survival over fecundity, independent of density effects (but see Bassar et al., 2013; N. L. Fowler et al., 2006, where all life history components played a comparable role in driving population growth under varying density). We hope that further studies will address the problem of why population growth rate is more sensitive to DD in survival.

We now discuss some limitations of our study. First, we pooled all life stages when counting per capita offspring, likely overestimating the number of reproductive adults that could drive subsequent population growth. Although mortality after pupation is very low, mortality at the larval stage is not negligible. Thus, we have limited ability to extrapolate our single-generation experimental outcomes across several generations. Second, because we could not explicitly account for cannibalism of eggs by larvae and adults, we likely underestimated fecundity and overestimated survival. Whether the strength of these effects changes with density remains unclear, and should be addressed in future work. Third, our quantification of the shape of DD (based on the difference in area between a nonlinear and linear model) is limited, in that more complex shapes (such as sigmoidal and quadratic) cannot be meaningfully quantified using this approach. However, for the large number of organisms for which the concave/convex/linear distinction is valid (Sibly et al., 2005; Thibaut & Connolly, 2020), this approach should work, with the benefit that it allows comparison between many different models whose parameters cannot be easily directly compared. We also note that the values of slope and shape defined in this way would depend on the units in which density is measured. However, when comparing parameters to determine their impact on slope and shape, the relative effect sizes are still meaningful. Finally, the models we used were fully deterministic, and it remains to be seen whether our conclusions are robust to the incorporation of stochasticity.

To our knowledge, our work is the first study to generate experimental data for DD in fecundity and survival across multiple habitats and populations, allowing us to disentangle the effects of environment, population, and density on the demographic parameters affecting density-dependent growth rate. The simplicity of our model, encapsulating key demographic traits into survival and fecundity, means that our framework can be applied across a very wide range of taxa. Our results also have interesting implications for the evolution of density dependence. For instance, we predict that any trait change that increases parameter *K* (tolerance of proportion survival to density) should be the primary mechanism of evolution under density dependent selection (e.g., Venkitachalam et al., 2022). Broadly, we also suggest that any factors that affect DD in survival are likely to have a strong effect on population dynamics. Therefore, population management strategies, e.g., for pest species, should consider not just controlling the number of adults or their fecundity, but specifically focus on juvenile survival. Further, our observation that the form of DD in *T. castaneum* is primarily concave suggests that a severe reduction in density (e.g., by pest control measures) would lead to proportionally rapid population growth (e.g. Powell, 1988). Perhaps a strategy focused on reducing population growth rates (e.g., by decreasing the temperature, hence slowing development) may be more effective in controlling the population of this pest. In summary, a deeper understanding of the form of DD and the demographic traits underlying it may have several important and interesting implications for both basic and applied ecology.

## ACKNOWLEDGEMENTS

We thank Shubha Govindarajan and Sushmita Krishnan for help with experiments, Amitabh Joshi for his population dynamics course and discussions, Shivansh Singhal and Soumya Panyam for discussions, and Vrinda Ravi Kumar and Pratibha Sanjenbam for comments on the manuscript. We acknowledge support and funding from the National Centre for Biological Sciences (NCBS–TIFR) and the Department of Atomic Energy, Government of India (Project Identification No. RTI 4006).

