## Supporting Information for "Density dependent survival drives variation in density dependent population growth of an insect pest"

**SUPPLEMENTARY METHODS**

**Stock populations**

We used ten source populations of *Tribolium castaneum* in this study (denoted 1, 2, 6, 7, 11, 12, 13, 18, 20, 24), each wild-collected from different locations in India and maintained in large laboratory stock populations for 5-10 years in a dark incubator, at 34 °C, on whole wheat flour. We maintained population stocks on a 7-8 week discrete generation cycle with 4-7 days of egg laying to initiate each generation. We provided at least 500 adults with 750g fresh wheat flour at the start of the egg laying, and replaced the flour and removed dead individuals and larval moults after 3-4 weeks. We collected adults for experiments at 2 weeks post-eclosion from stock boxes (~55 days post egg hatching).

**Additional methods for Experiment 1**

We aimed to measure one replicate for every possible combination of habitat (4 flour habitats: wheat, sorghum, finger millet, and corn), population (10 source populations) and density (10 densities, ranging from 0.6 to 7 beetles per g flour) (total 400 combinations requiring a total of 1500 adults per population) in 2018. Because some stock populations did not have enough individuals, we could only set up 336 combinations. We therefore repeated part of this experiment in 2019, to better sample some of the population-habitat combinations across the density range. We measured 12 combinations of population and habitat across ten densities, and for combinations that were measured in both years (total 3 such combinations), we used only the second measure for data analysis. In the final dataset, we have 384 combinations, representing all 40 population-habitat combinations, and at least 9 (out of 10 planned) densities for each population-habitat combination (except for population 6 in wheat flour, which had only 6 densities, but this range included the lowest and highest densities).

**Methods for Experiment 3**

Here, using four habitats, four populations, two densities (1.2 and 6 beetles per gram flour in 50g, which is 60 and 300 beetles respectively), and three replicates per combination of habitat, population, and density, we tested whether DD in per capita offspring varies across habitats and populations. This enabled us to overcome the previous limitation of low replication in Experiment 1. Note that the methods for collecting the adults were the same as in Experiment 1, except the collected adults were about two weeks older than in Experiment 1 due to logistical constraints. We set up the densities for one week in double sifted flour and removed adults along with 10g of the flour after one week. The remaining 40g was left undisturbed to allow offspring development for 3 weeks post adult removal (wheat and sorghum) or 4 weeks post adult removal (corn and finger millet). The changes to the experimental protocol (double sifting and removal of 10g flour) allowed us to sample the number of eggs, the data for which is not presented here. To estimate the total number of offspring, we multiplied the counted number of offspring (from 40 g flour) by 1.25, to scale up to 50 grams of total flour.

We used an ANOVA to determine the effect of habitat, population and density on per capita offspring (Table S2; Figure S3). To increase homoscedasticity of the data, we square root transformed per capita offspring. We tested the impact of each potential explanatory variable (source population, habitat, and density) as well as their interaction effects.

**Methods for Experiment 4**

In Experiment 2, we did not sample low egg densities, so we could not experimentally validate the intercept of proportion survival at a value of 1. Hence, we separately estimated egg survival at very low egg densities, where there would be no competition. We collected eggs from stocks of population 12, and placed 1 egg in 1g of flour, in 1.5 ml Eppendorf tubes (n = 96-100 eggs for each habitat). We counted surviving individuals after 25 days (wheat and sorghum) or 32 days (corn and finger millet). The proportion survival was close to 1 for all habitats (varies from 0.83-0.96, SI data), and did not differ significantly across habitats (Table S3).

**SUPPLEMENTARY TABLES**

**Table S1**: Parameter values and 95% confidence intervals (CI) for the fecundity and survival relationships for population 12 in each habitat (Experiment 2). Curves were fit using nonlinear least squares. Per capita fecundity = *a**density*^b^.* Proportion survival = (1+*alpha**(total eggs/*K*))/(1+(total eggs/*K*)). W = Wheat flour, C = Corn flour, FM = Finger millet flour, S = Sorghum flour.

| **Habitat** | **Fecundity *a*** | **Fecundity *b*** | **Survival *alpha*** | **Survival *K*** |
| --- | --- | --- | --- | --- |
| W | 140.5 (71.3) | -0.45 (0.20) | 0.010 (0.049) | 127.4 (35.7) |
| C | 70.5 (49.3) | -0.47(0.26) | 0.171 (0.052) | 13.7 (16.0) |
| FM | 144.7(75.1) | -0.57 (0.21) | 0.058 (0.046) | 34.4 (21.4) |
| S | 146.2 (63.8) | -0.52 (0.17) | 0.062 (0.120) | 188.0 (97.0) |

**Table S2:** Results of an ANOVA performed for factors affecting the square root of per capita offspring (Experiment 3). Per capita offspring ~ density*habitat*population (adjusted r squared = 0.92). Terms with significant effects are in bold. “pop” stands for population.

| **Per capita offspring as explained by:** | **Degrees of freedom** | **Sum of squares** | **Mean squares** | **F value** | **P value** |
| --- | --- | --- | --- | --- | --- |
| **density** | **1** | **120.934315** | **120.934315** | **1015.10014** | **5.62E-41** |
| **habitat** | **3** | **10.2061826** | **3.40206086** | **28.5562658** | **7.71E-12** |
| **pop** | **3** | **3.10389699** | **1.03463233** | **8.68451128** | **6.45E-05** |
| **density:habitat** | **3** | **2.85980581** | **0.9532686** | **8.0015593** | **0.00013126** |
| density:pop | 3 | 0.56224856 | 0.18741619 | 1.57313659 | 0.20451839 |
| habitat:pop | 9 | 1.61626891 | 0.17958543 | 1.50740672 | 0.16454946 |
| density:habitat:pop | 9 | 1.50291702 | 0.16699078 | 1.40168954 | 0.20629352 |
| Residuals | 64 | 7.6246627 | 0.11913535 | NA | NA |

**Table S3:** Results of a binomial GLM performed for the effect of habitat on the probability of survival of isolated eggs (n=100, except sorghum, where n= 96; Experiment 4). Probabilities of survival are given in SI Data, Experiment 4.

| **Probability of egg survival explained by:** | **estimate** | **std.error** | **statistic** | **p.value** |
| --- | --- | --- | --- | --- |
| (Intercept) | 2.197 | 0.333 | 6.592 | 4.35E-11 |
| Sorghum | -0.588 | 0.431 | -1.362 | 0.1730 |
| Finger millet | 0.981 | 0.610 | 1.609 | 0.108 |
| Wheat | 0.981 | 0.610 | 1.609 | 0.108 |

**SUPPLEMENTARY FIGURES**

**Figure S1: Schematic explaining our simulations to analyse the impact of variation in parameters of DD in different demographic traits on DD in population growth rate.** This analysis was conducted for data derived from Experiment 2, separately for both populations (populations 7 and 12). Number of simulated values of the different parameters in population 7 and 12, respectively: fecundity *a*: 163, 152; fecundity *b*: 364, 114; survival *alpha*: 62, 161; survival *K*: 97, 175. Total number of simulations for population 7 and 12 (calculated by summing the above values and multiplying by 4, for all habitats) = 3104 and 2408 respectively.

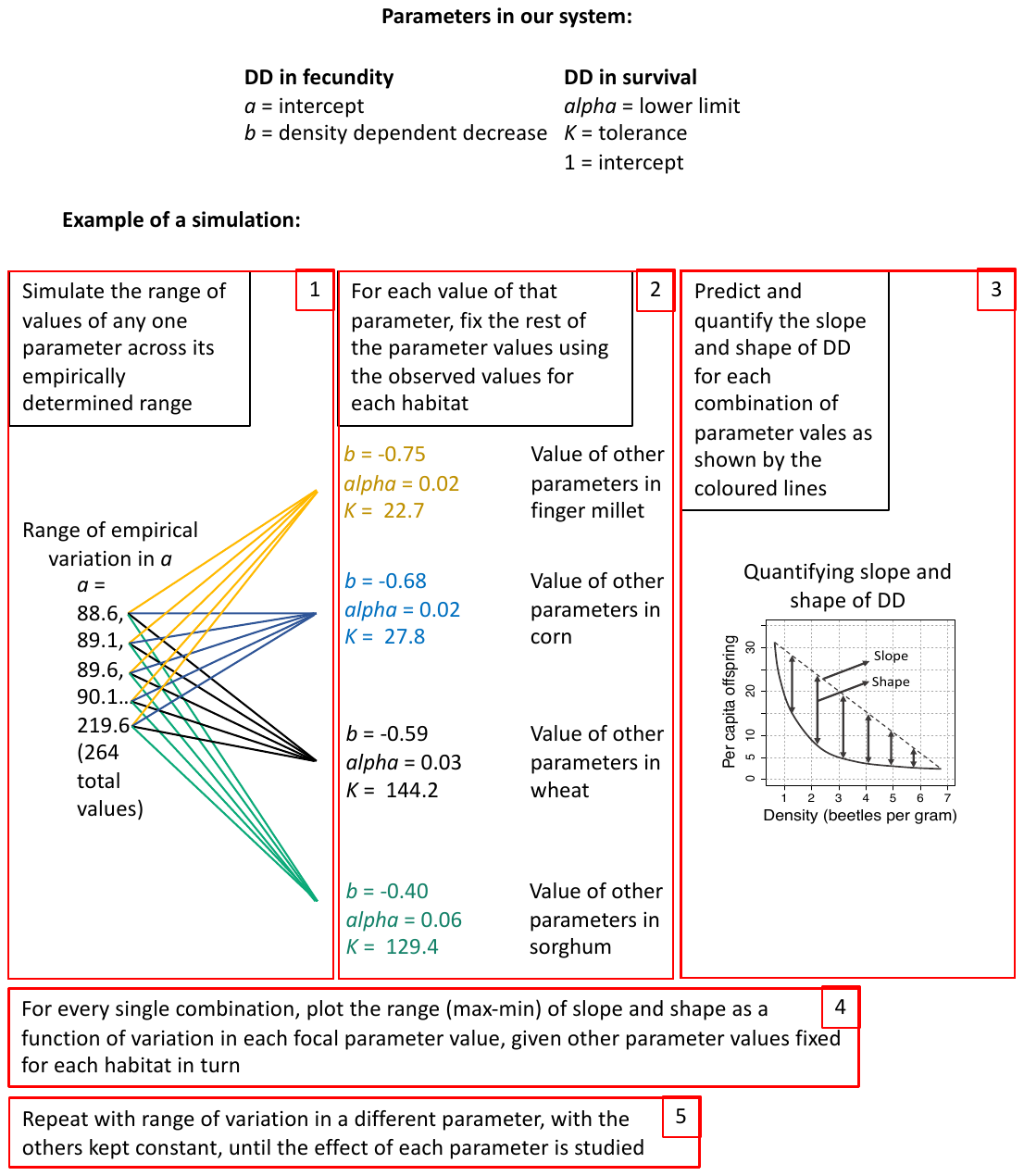

**Figure S2: DD in fecundity and survival accurately predicts DD in per capita offspring.** The figure shows data for population 12 measured in Experiment 2; results for population 12 are shown in Figure 3. (**A–C**) Density dependence in per capita fecundity (as a function of adult density), in survival (as a function of number of eggs), and in per capita offspring (as a function of adult density) in wheat flour. Each point represents experimentally measured data for a single replicate (n = 2–5 per density per habitat). In panels **A** and **B**, dashed lines indicate models fitted to the data using nonlinear least squares. In panel **C**, the solid line shows the predicted density dependence in per capita offspring derived from multiplying the equations for models shown in panels **A** and **B**. Similarly, panels **D–F** show results for finger millet, **G–I** show results for corn, and **J–L** show results for sorghum. Model equations and estimated parameter values are shown in Table S1 and its legend.

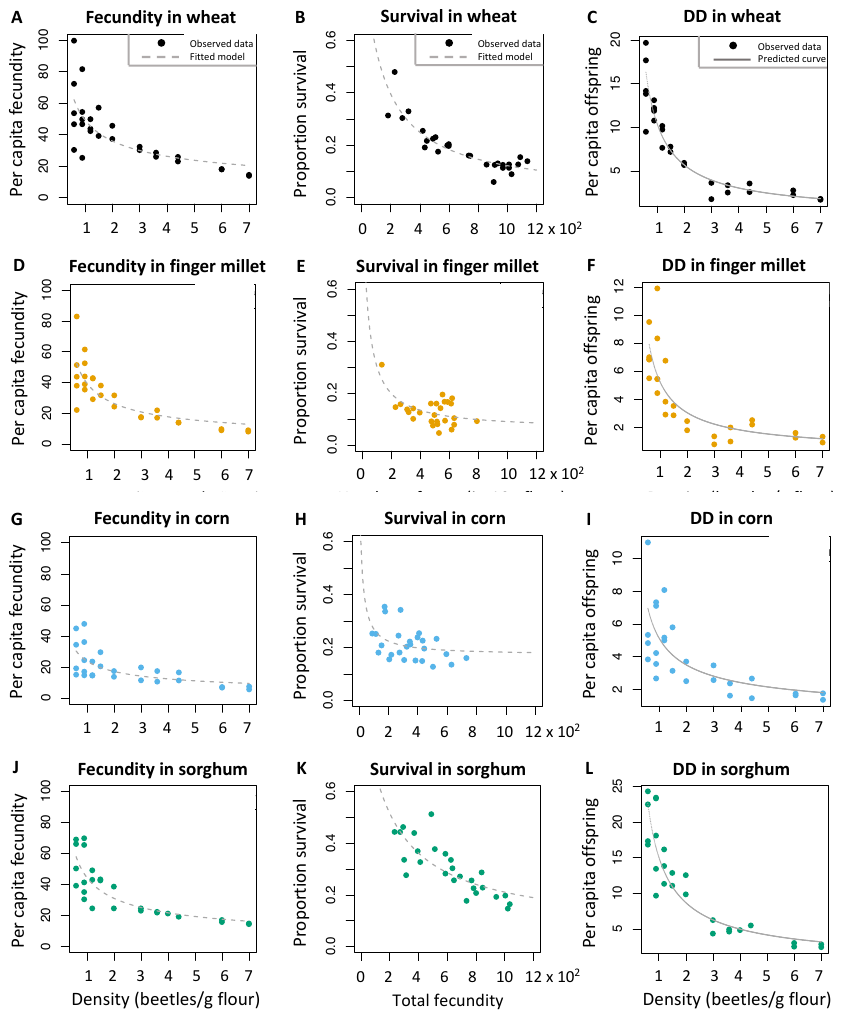

**Figure S3: DD varies across habitats but not populations in experiment 3.** Using data from 4 source populations at 2 densities in 4 habitats, and 3 replicates for each combination (Experiment 3), we analysed how DD in the square root of per capita offspring varies across habitats and populations (Table S2). Here, we plot the main result, that DD in square root of per capita offspring varies across habitats but not populations. In panel **A**, the data are organised by population, while in panel **B**, they are organised by habitat. Error bars show ±95% confidence intervals.

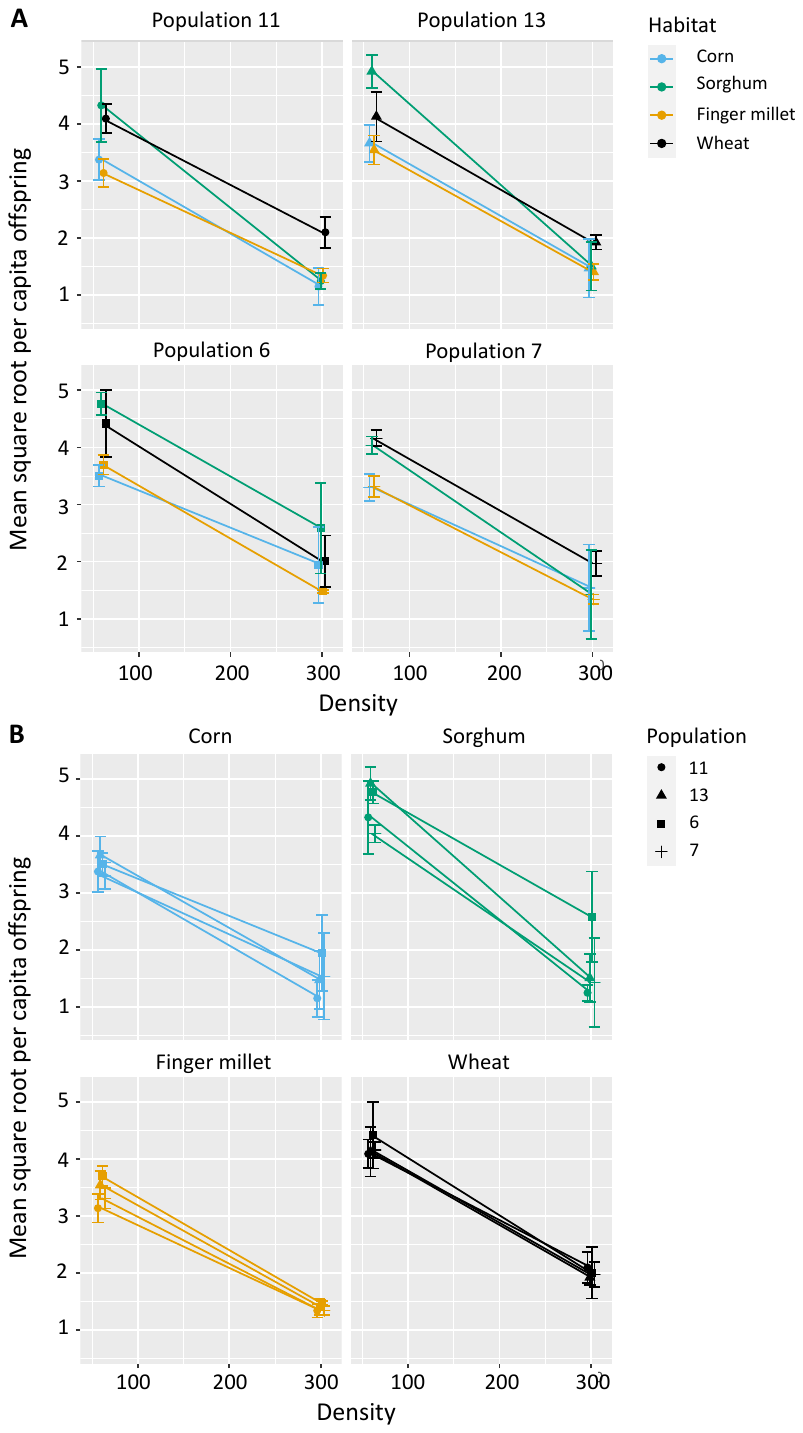

**Figure S4: The slope and shape of DD in per capita offspring are most sensitive to survival parameter *K.*** Plots show results of simulations to test the effect of varying each parameter value (Table S1) in our models describing DD in fecundity and survival for population 12 (Figure S2). Results for population 7 are shown in Figure 4. (**A–B**) For each habitat we varied the value of one parameter in turn, keeping other parameters constant. Plots show the range of the resulting values of the slope and shape of DD in per capita offspring, predicted as described in Figure 3 (across 3104 simulations). (**C–D**) Here, we again simulated variation in a single parameter value for a given habitat, but only increased the parameter value by 1% of the observed value. Plots show the difference in the resulting slope and shape of DD, from observed values. Points are slightly jittered about the x axis for clarity. Details of the simulations are given in the Methods section and in Figure S1.

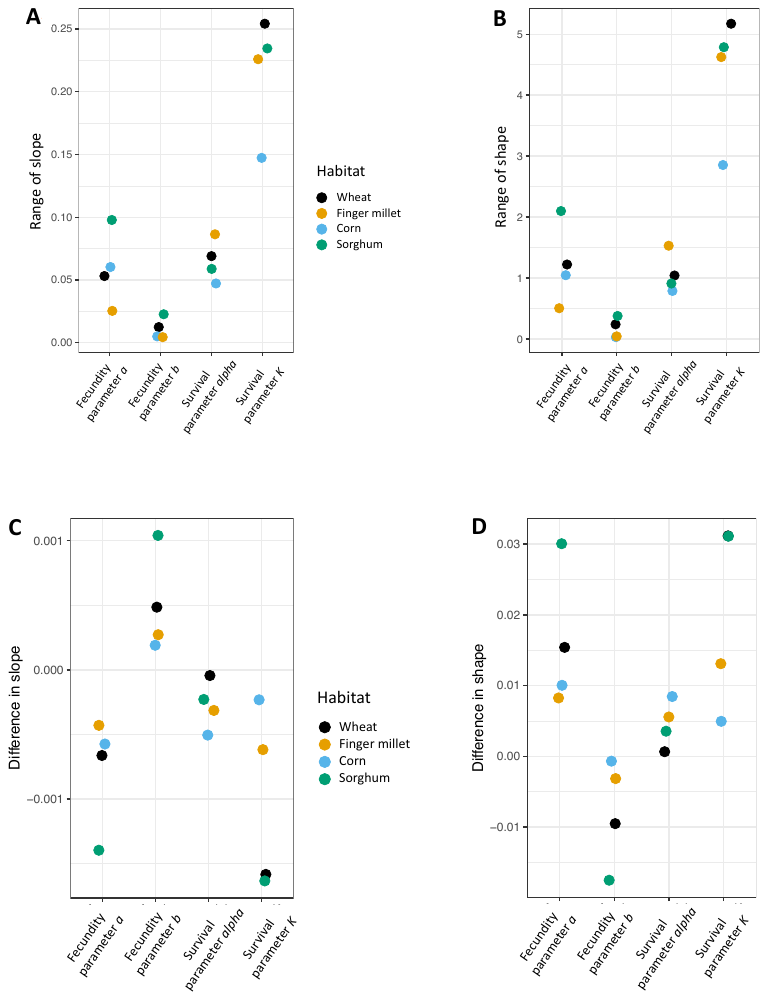

**Figure S5: The impact of proportional variation in survival parameter *K* is robust to how we calculate its effect.** For models fit to data for population 7 from Experiment 2, we varied each parameter by multiplying it with 1.01, as described for population 12 above (Figure S4 C–D) and for population 7 in the main text (Figure 4C-D). However, instead of calculating the difference between the resulting slope (**A**) and shape (**B**) of DD and observed slope or shape, we estimated the effect of altering parameter values using a ratio (simulated/observed; a value of 1 on the y axis indicates no change). The trends are similar in both cases (compare this Figure with Figure 4C-D and S4C-D), suggesting that our results are robust to the method of calculation. Points are slightly jittered about the x axis for clarity.

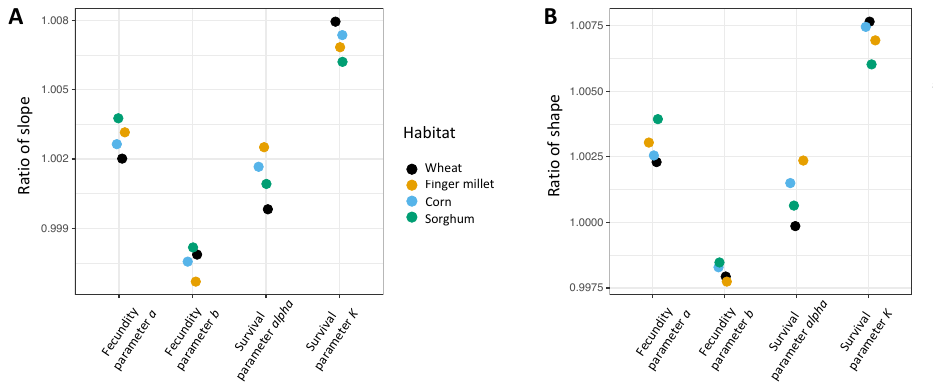

**Figure S6: The slope and shape of DD in per capita offspring are most sensitive to survival parameter *K* when varied proportionally by up to 100%.** As described in Figures S1 and S4, we varied parameter values for population 7 by 5%, 20%, 50% or 100%, this time in both directions. (**A–H**) Change in slope and shape of DD population growth after increasing parameter values proportionally. (**I–N**) Change in slope and shape after decreasing parameter values proportionally. Empirically, the minimum variation in parameter values across habitats was 100% (Table 1: fecundity *b* ranges from -0.4 to -0.75). Simulating >100% changes in parameter values would alter some parameters far beyond the empirically realistic range, so we did not simulate larger changes. We did not implement a 100% decrease as that would bring the parameter value to zero. We found largely similar results as observed with 1% parameter changes, i.e., a larger impact of proportional variation in survival *K* than fecundity *a*. However, the difference between the impact of these two parameters is slightly smaller when the parameters were decreased rather than increased proportionally; e.g., compare panels A and B with panels I and J (note different axes ranges).

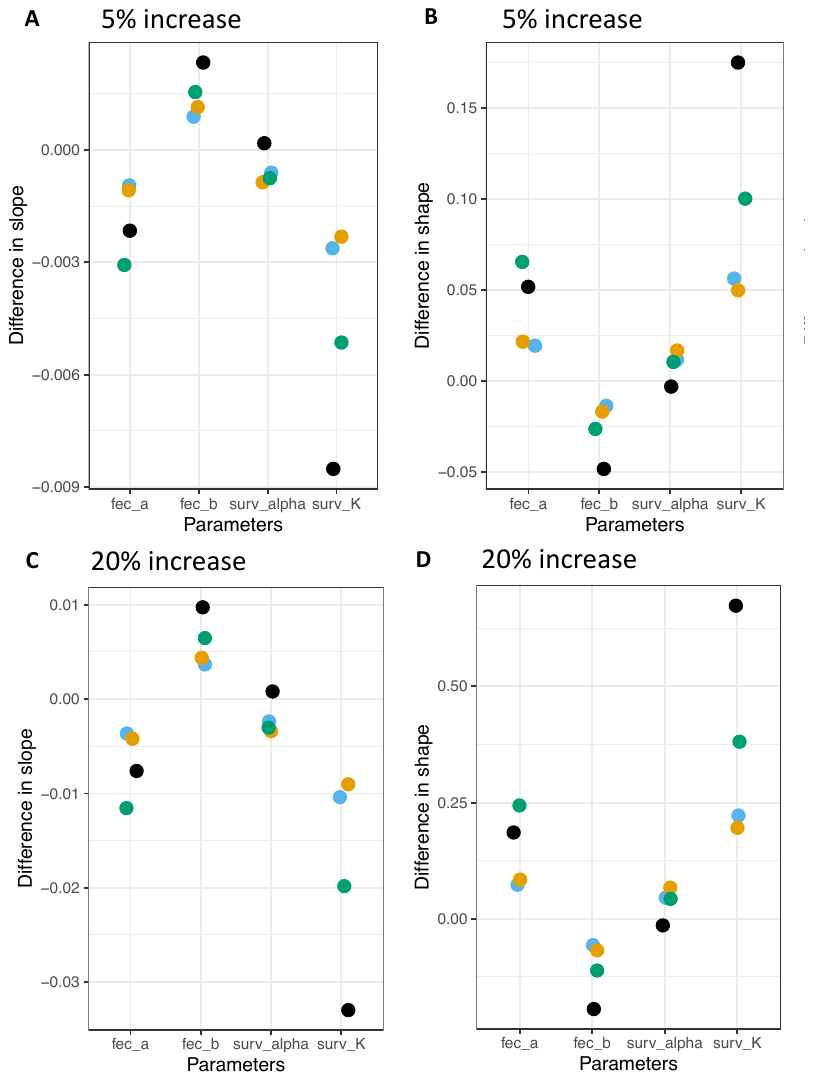

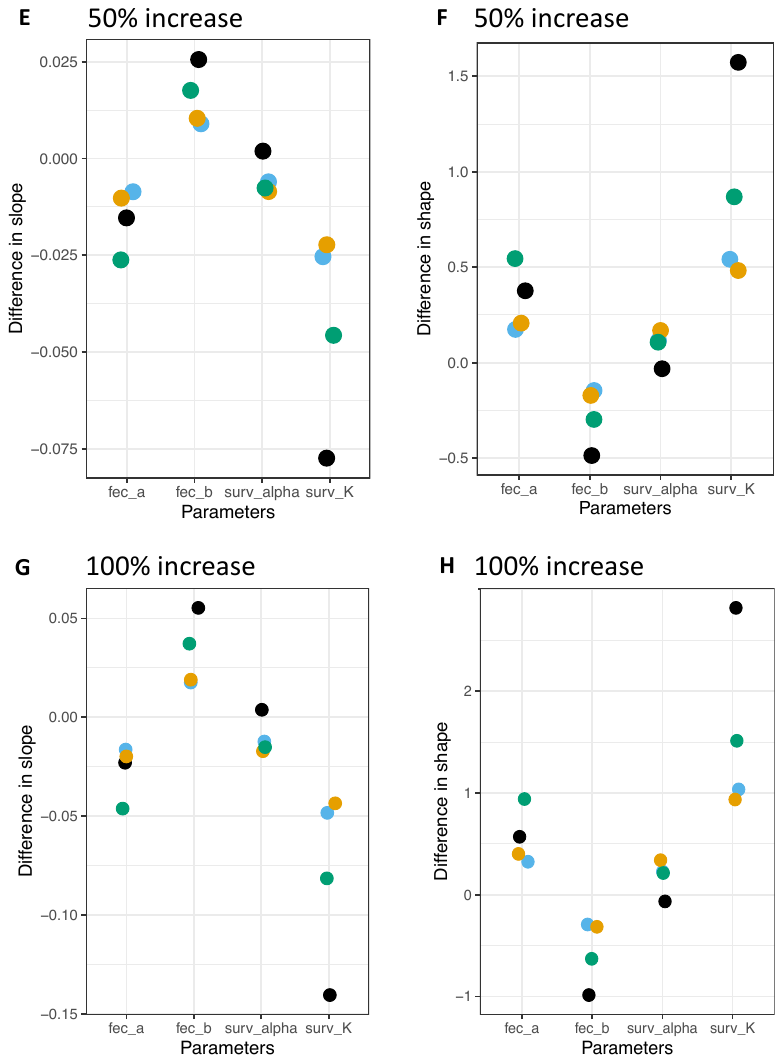

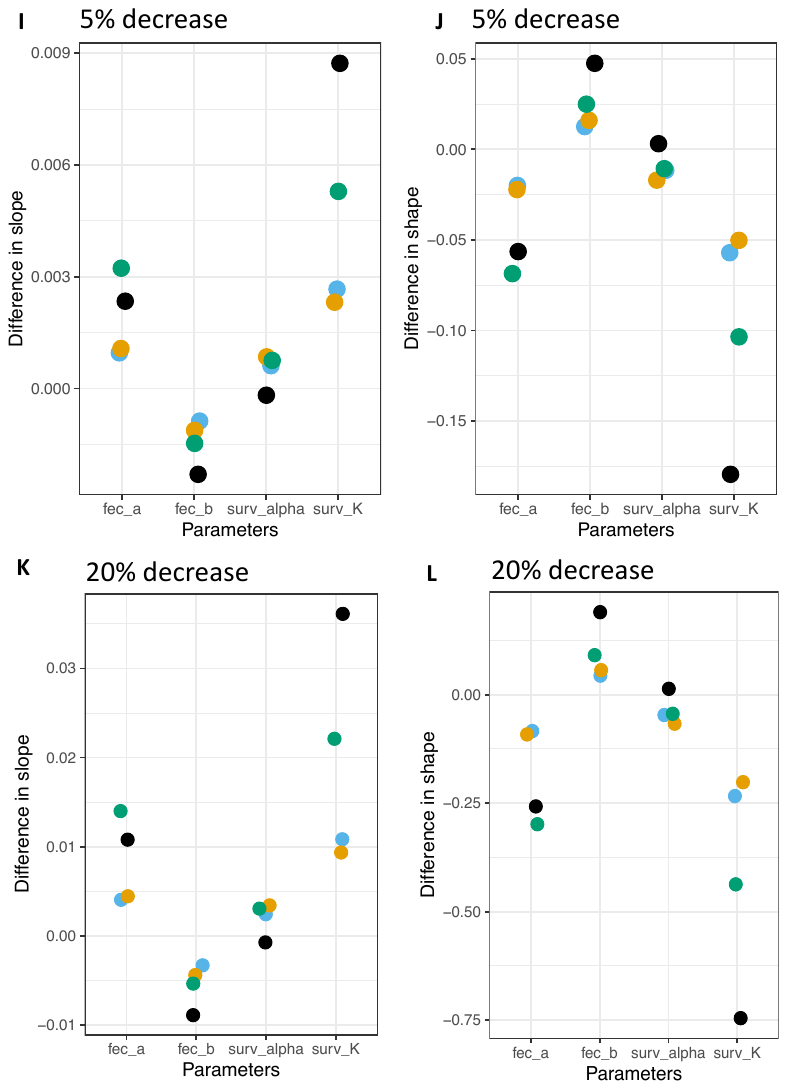

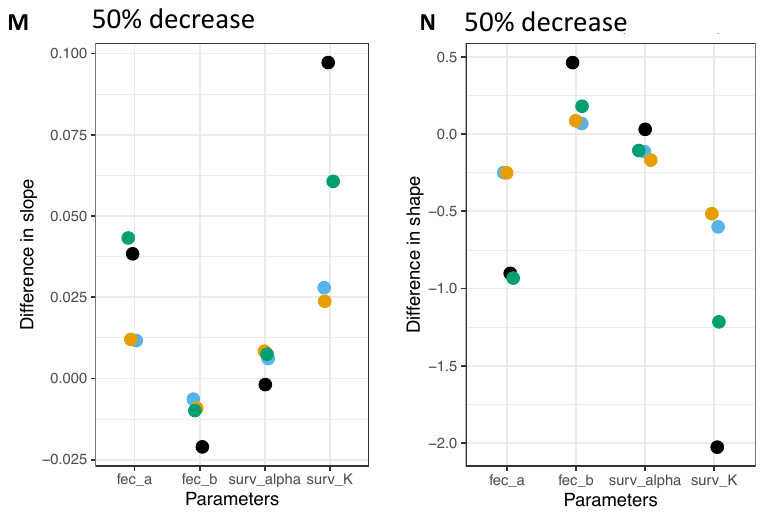

**Figure S7: The slope and shape of DD in per capita offspring are most sensitive to survival parameter *K* and fecundity parameter *a* when varied proportionally up to 20%.** As in Figure S6, we varied parameter values for population 12 but only by 5% and 20%. (**A–D**) Change in slope and shape of DD population growth after increasing parameter values proportionally. (**E–H**) Change in slope and shape after decreasing parameter values proportionally. For population 12, the least variation in empirically estimated parameter values was 25% (Table S1: fecundity *b* ranges from -0.45 to -0.57), so we only tested the effect of 5% and 20% change in parameter values. We found primarily the same results as due to 1% variation, i.e., an equal impact of proportional variation in survival *K* as fecundity *a*.

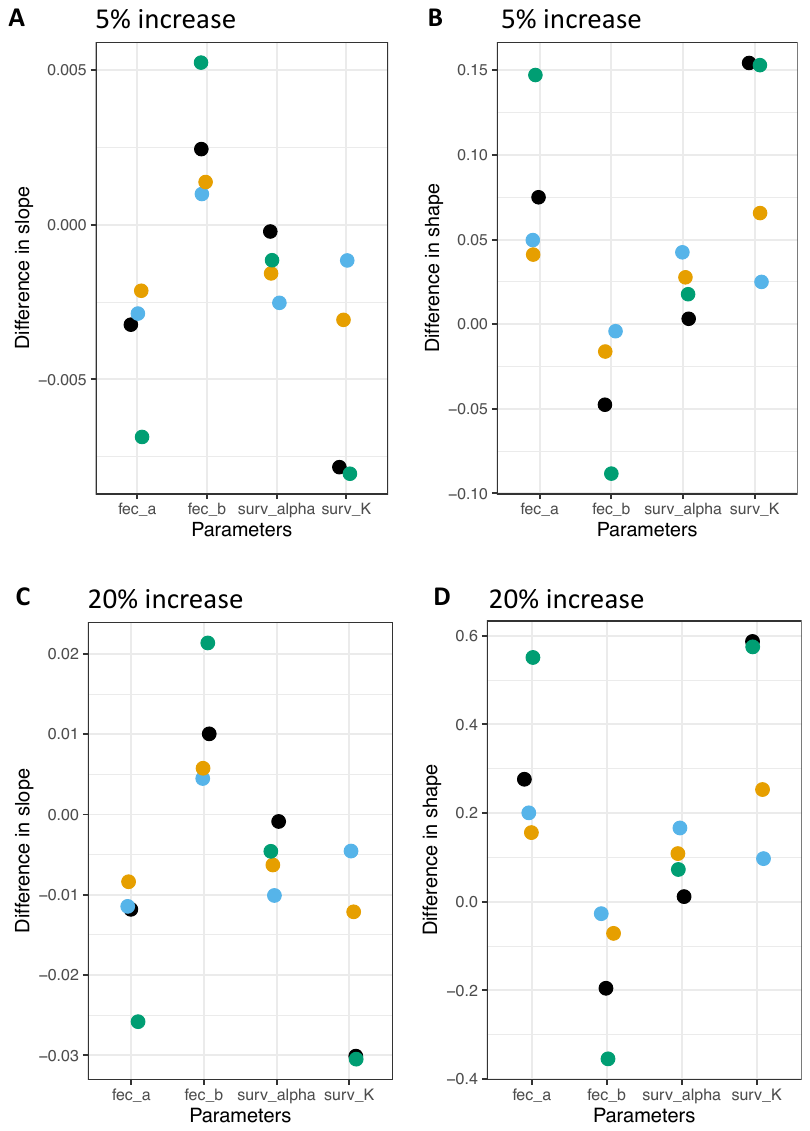

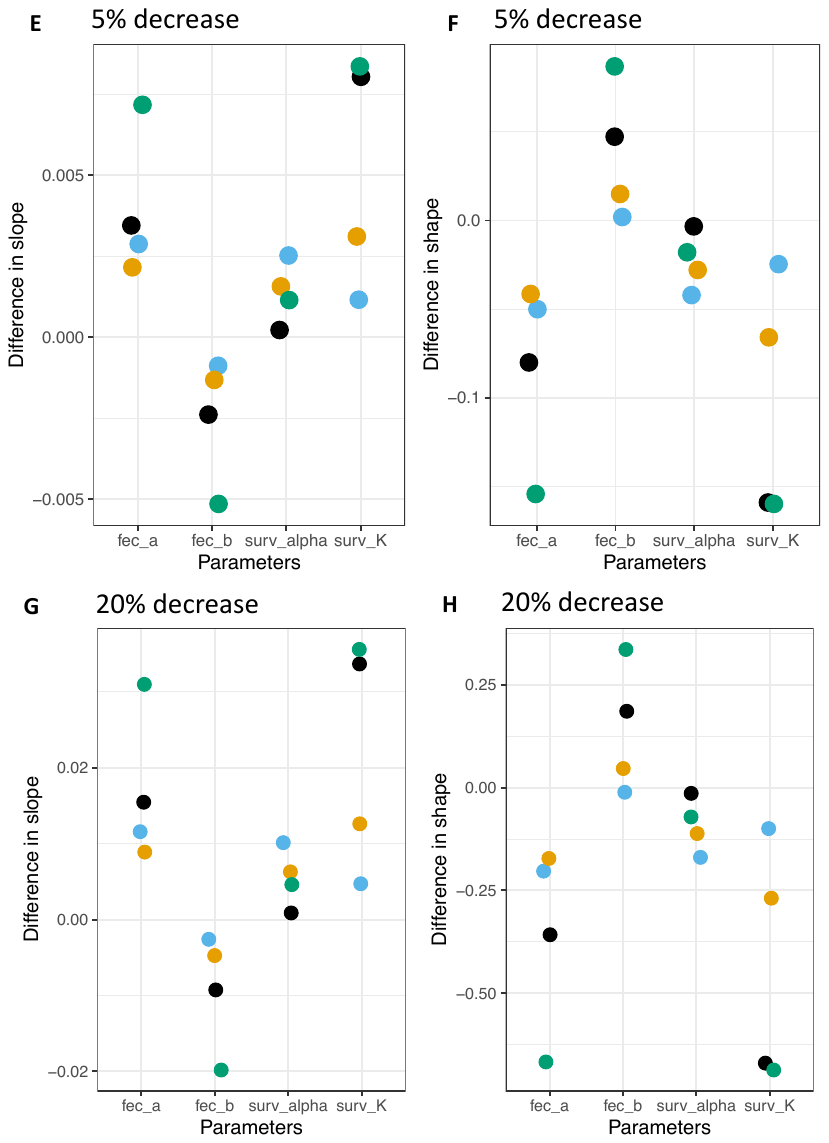
